# Imprinted maternally-expressed microRNAs antagonize paternally-driven gene programs in neurons

**DOI:** 10.1101/717868

**Authors:** Amanda J. Whipple, Hannah N. Jacobs, Vincent Breton-Provencher, Mriganka Sur, Phillip A. Sharp

## Abstract

Imprinted genes with parental-biased expression are hypothesized to result from an evolutionary conflict between the parental genomes over procurement of maternal resources. Accordingly, imprinted genes are enriched in pathways regulating nutrient acquisition, energy homeostasis, and growth. Here, we functionally characterize a large cluster of maternally-expressed microRNAs (miRNAs) to explore why they evolved imprinted expression in neurons. Using an induced neuron (iN) culture system, we show maternally-expressed miRNAs from the miR-379/410 cluster repress paternally-expressed genes, including known regulators of energy homeostasis *Plagl1* and *Peg3*. Additional non-imprinted metabolic regulators are also co-targeted by miR-379/410. Maternal deletion of this imprinted miRNA cluster results in de-repression of its targets and up-regulation of a broader gene program regulating feeding behavior and synaptic transmission. These data suggest non-coding RNAs actively engage in parental genomic conflict, whereby maternally-expressed miRNAs antagonize paternally-driven gene programs in neurons.

## Introduction

Genomic imprinting is an epigenetic process that results in preferential gene expression from the maternal or paternal allele. Imprinting negates the ability of diploid organisms to buffer against recessive mutations, and the evolutionary pressures driving imprinting are openly debated (Haig, 2014; Patten et al., 2014; Spencer and Clark, 2014). One widely-cited theory, known as the parental conflict or kinship theory, posits that the two parental genomes within offspring have opposing interests in the distribution of maternal resources (Haig and Westoby, 1989; Moore and Haig, 1991). Paternally-expressed genes, such as Insulin-like growth factor 2 (*Igf2*), tend to stimulate nutrient uptake and drive embryonic growth (DeChiara et al., 1991; Ferguson-Smith et al., 1991). In contrast, maternally-expressed genes within offspring negatively regulate growth, thereby conserving maternal resources. An example is the maternally-expressed *Igf2* receptor, which is a direct functional antagonist of *Igf2* signaling in the placenta (Barlow et al., 1991). In accordance with this theory, imprinted genes are enriched for diverse metabolic functions and may serve to coordinate growth with energy homeostasis (Peters, 2014). Many microRNAs (miRNAs) have evolved imprinted expression, but the potential evolutionary benefits of parental-biased miRNA expression remain to be determined.

miRNAs direct post-transcriptional repression of mRNA targets through the RNA induced silencing complex (RISC). There are three recently evolved, imprinted miRNA clusters. miRNA clusters in the eutherian-specific *Maternally-expressed gene 3* (*Meg3)* are maternally-expressed in multiple tissues including the brain (Seitz et al., 2003), whereas the rodent-specific *Sfmbt2* and primate-specific *C19MC* miRNA clusters are paternally-expressed in the placenta (Noguer-Dance et al., 2010; Wang et al., 2011). While miRNAs typically exert modest effects on gene expression, the physiological effects of miRNA clusters may be amplified by coordinated co-expression of multiple miRNAs and co-targeting of multiple mRNAs in a pathway. Interestingly, a key RISC component, *Argonaute2* (*Ago2)*, also exhibits maternally-biased expression in the brain (Andergassen et al., 2017; Bonthuis et al., 2015; Perez et al., 2015), suggesting the maternal genome may benefit from miRNA activity in the brain. Therefore, we sought to characterize the potential cellular and physiological impact of maternally-expressed miRNAs in neurons.

A dense cluster of 39 miRNAs, miR-379/410, is located in the terminal region of *Meg3* known as *Mirg* (Figure 1A). The miR-379/410 cluster is expressed during early development and in postnatal brain (Labialle et al., 2014). Maternal inheritance of miR-379/410 deletion resulted in hypoglycemia immediately after birth (Labialle et al., 2014), suggesting these miRNAs control metabolic physiology during early development. Adult mice with maternal miR-379/410 deletion display anxiety-related behaviors and increased sociability, perhaps due to alterations in synaptic signaling (Lackinger et al., 2018; Marty et al., 2016). One miRNA from the cluster, miR-134-5p, localizes to the synapse and regulates dendritic spine size through translational repression of *Limk1* and *Pum2* (Fiore et al., 2009; Schratt et al., 2006).However, relatively few targets of this large miRNA cluster have been identified, and molecular understanding of how these miRNAs may coordinately regulate cellular programs is lacking.

**Figure 1.**
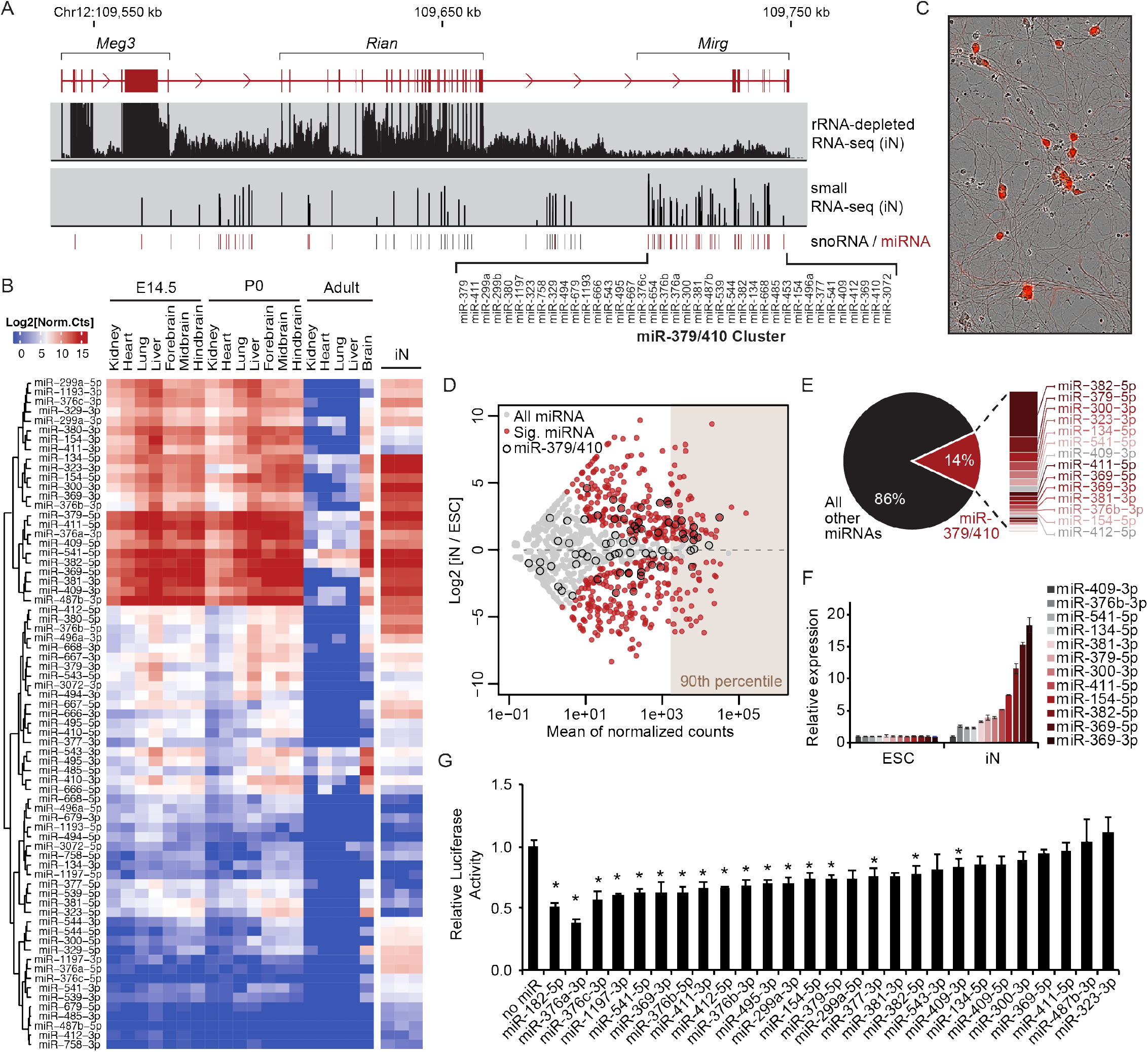
miR-379/410 miRNAs are active in induced neurons. **A.** Schematic of the annotated *Meg3* long non-coding RNA and small non-coding RNAs processed from the locus (snoRNA, small nucleolar RNA; miRNA). The location of the miR-379/410 cluster is indicated, along with genomic alignment of rRNA-depleted RNA-seq and small RNA-seq data from iNs. **B.** Heatmap of normalized counts of mature miRNAs from the miR-379/410 cluster generated by small RNA-seq of mouse tissues at embryonic day 14.5 (E14.5), postnatal day 0 (P0), adult tissues, and *in vitro* differentiated iNs (n = 3). Rows are organized by unsupervised hierarchical clustering. **C.** Overlay of brightfield and Ngn2-mCherry fluorescence image of iNs. **D.** Expression changes of miRNAs upon differentiation shown as an MA plot. Gray, all miRNAs; Red, differentially expressed miRNAs (p ≤ 0.05, fold change ≥ 2); Black outline, miR-379/410. miRNAs in the 90^th^ percentile of expression are indicated. **E.** Pie chart indicating the proportion of small RNA-seq reads mapping to miR-379/410 miRNAs versus other miRNAs in iNs. **F.** qRT-PCR for mature miRNAs from miR-379/410 in iNs relative to ESCs. Mean ± SEM (n = 3). **G.** Bar graph of Firefly luciferase activity with or without 3’ UTR miRNA target sites relative to Renilla luciferase internal control. Mean ± SEM (n = 6); *p ≤ 0.05, calculated using unpaired two-tailed Student’s t-test. See also Figure S1.

In this study, we sought to understand why miRNAs evolved imprinted expression by performing a comprehensive characterization of miR-379/410 targets in neurons. We generated a powerful discovery model for imprinted miRNA activity using Neurogenin2-induced differentiation of hybrid mouse embryonic stem cells (ESCs). Using this system, we identified 33 active miRNAs from the imprinted miR-379/410 cluster. We found that these maternally-expressed miRNAs direct the RISC machinery to paternally-expressed genes, which are de-repressed upon maternal miR-379/410 deletion. We also identified additional miR-379/410 targets that function in metabolic regulation. Moreover, we determined that neurons with maternal miR-379/410 deletion have aberrant up-regulation of genes associated with feeding behavior and synaptic transmission, indicating miR-379/410 may function in energy homeostasis. These data suggest the maternal genome utilizes the miRNA pathway to antagonistically regulate paternally-driven gene programs in neurons.

## Results

### miR-379/410 expression is induced upon neuron differentiation

To determine the relative levels of mature miRNAs produced from the miR-379/410 cluster during development, we analyzed small RNA-seq data from mouse peripheral tissues and brain regions at embryonic, early postnatal, and adult developmental stages (Figure 1B). A distinct subgroup of 24 mature miRNAs located throughout the cluster are highly expressed across all embryonic and early postnatal tissues. These miRNAs are down-regulated during postnatal development resulting in little to no expression of miR-379/410 in adult tissues except brain, which maintains high expression in adulthood. The *Meg3*/*Mirg* host gene is also abundant in the brain (Figure S1A) and shows higher expression in neurons than astrocytes, microglia, or other brain-derived cell types (Figure S1B).

We established an *in vitro* neuron differentiation system to study the maternally-expressed miR-379/410 cluster. Doxycycline-inducible *Neurogenin2* (*Ngn2*) was stably integrated into F1 hybrid mouse ESCs from a *Mus. musculus* × *Mus. castaneous* cross (Eggan et al., 2001) that allows maternal and paternal alleles to be distinguished by single nucleotide polymorphisms (SNPs) (Figure 1C, Figure S1C). Upon differentiation by doxycycline treatment, induced neurons (iNs) expressed markers of glutamatergic neurons and displayed spontaneous electrophysiological activity (Figure S1D-G). We performed small RNA-seq to determine the dynamics of miRNA expression upon neuron differentiation. We found that miR-379/410 expression in iNs mimics *in vivo* expression (Figure 1B). Many miRNAs from the cluster are amongst the most highly expressed miRNAs upon differentiation (Figure 1D). In fact, we found that 14% of all miRNAs detected in iNs were from the miR-379/410 cluster (Figure 1E). We confirmed up-regulation of miR-379/410 in iNs by qRT-PCR and Northern blot (Figure 1F, Figure S1H). We conclude that a subset of the miR-379/410 cluster is highly expressed and dynamically regulated during development and differentiation.

We then asked if miR-379/410 showed repressive activity in iNs. Perfect target sites for individual miRNAs from the cluster were inserted into the 3’ UTR of a luciferase reporter, and luciferase activity was measured in iNs. We found sixteen miRNAs from the cluster significantly repressed luciferase activity by 10-60% (p ≤ 0.05) (Figure 1G). The perfect target site for miR-182-5p, an abundant miRNA in iNs, was used as a positive control and resulted in 50% repression of luciferase activity. These data indicate that Ngn2-induced neurons are a great system for identifying miR-379/410 targets.

### Identification of RISC-associated miRNAs in induced neurons

In order to determine which miRNAs and mRNA target sites are bound by the RISC complex in iNs, we performed Ago2 single-end enhanced crosslinking and immunoprecipitation (seCLIP-seq). We stably integrated a transgene expressing untagged-Ago2 or HA-Ago2 into ESCs, differentiated cells to iNs, and performed seCLIP-seq using HA antibody (Figure S2A). We identified 211 Ago2-bound miRNAs, including 27 miRNAs from the miR-379/410 cluster (p ≤ 0.01, fold change ≥ 2) (Figure 2A, Figure S2B). The miRNAs most highly bound by Ago2 were miR-300-3p, miR-541-5p, miR-485-3p, and miR-758-3p, which was independently confirmed by RNA immunoprecipitation followed by qRT-PCR (Figure S2C). The expression, reporter activity, and Ago2 association of all mature miRNAs from the miR-379/410 cluster are summarized in Figure S2D. Based on these data, we sought to identify targets of active miR-379/410 miRNAs, which we defined as the 33 miRNAs with significant Ago2 binding or repressive reporter activity.

**Figure 2.**
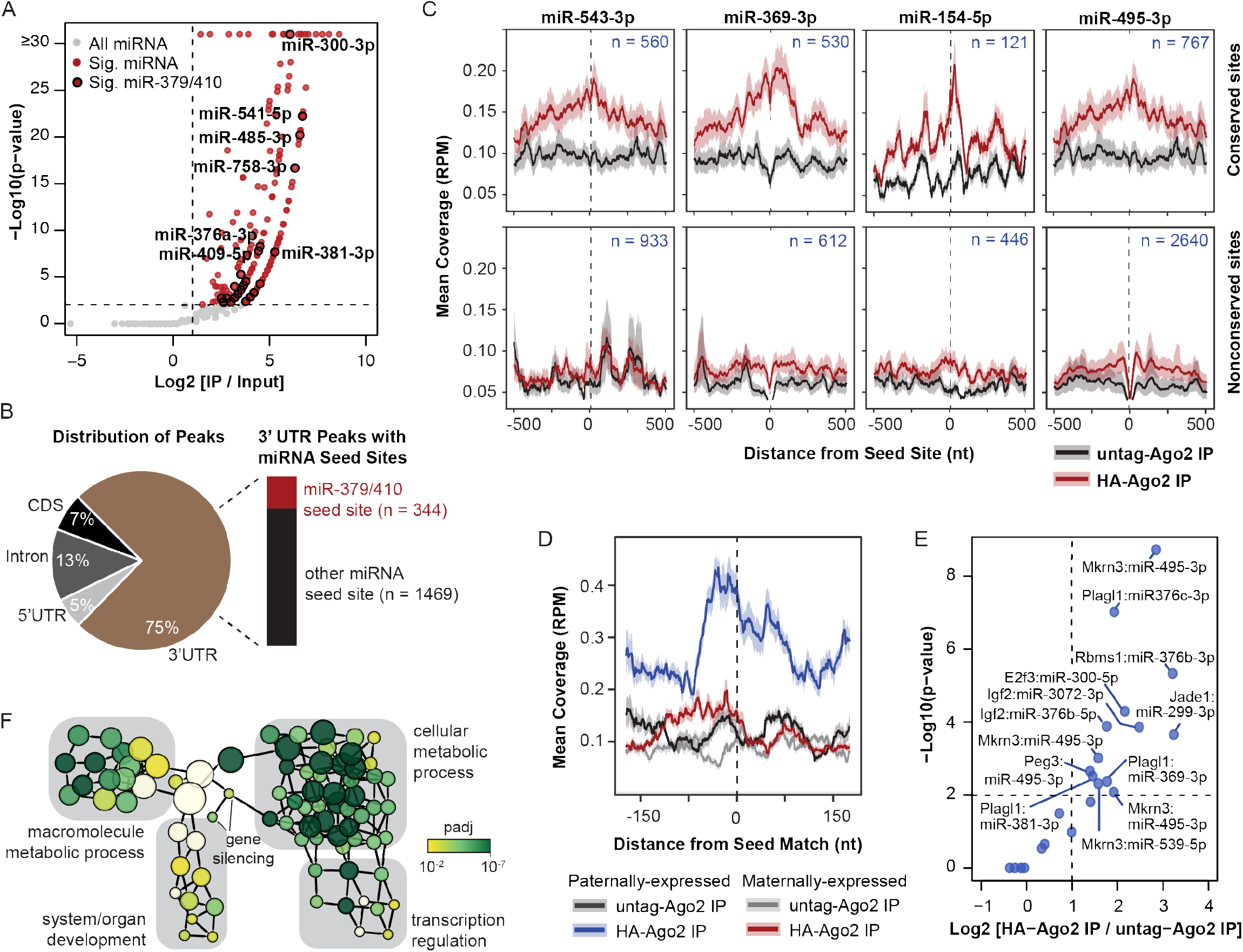
miR-379/410 directs Ago2 to paternally-expressed genes and other metabolic regulators. **A.** Volcano plot of seCLIP-seq reads mapped to miRNAs. Gray, all miRNAs (n = 2,045); Red, significantly enriched miRNAs (p ≤ 0.01, fold change ≥ 2, n = 211); Black outline, significantly enriched miR-379/410 (p ≤ 0.01, fold change ≥ 2, n = 27). The identities of the most enriched miR-379/410 miRNAs are labelled. **B.** Left: Pie chart indicating distribution of significant Ago2 peaks in 5’ UTR, Intron, CDS (coding sequence), and 3’ UTR. Right: Bar graph indicating the number of significant Ago2 peaks in 3’ UTRs that overlap with a seed site for an active miR-379/410 miRNA. **C.** Metagene plot of Ago2 seCLIP-seq coverage (reads per million, RPM) at seed sites for the indicated miRNAs, including coverage 500 nt upstream and downstream of the seed site. The number of sites included in each plot is shown. Top, conserved sites; Bottom, nonconserved sites with TargetScan score ≥ 90; Black, untagged-Ago2 IP; Red, HA-Ago2 IP. **D.** Metagene plot of Ago2 seCLIP-seq coverage (reads per million, RPM) at conserved seed sites for the active miR-379/410 in paternally-expressed genes (n = 38 seed sites) and maternally-expressed genes (n = 33 seed sites), including coverage 175 nt upstream and downstream of the seed site. **E.** Volcano plot of Ago2 peaks in paternally-expressed genes. Dashed lines, cutoffs for significant peaks (p ≤ 0.01, fold change ≥ 2). **F.** Biological network of gene ontology terms associated with miR-379/410 targets identified by Ago2 seCLIP-seq. The node size corresponds to the number of genes in the node. The node color indicates the adjusted p-value for the association of the gene ontology term with the miR-379/410 targets relative to the background of all expressed genes. See also Figure S2.

We called peaks in mapped seCLIP-seq reads that were significantly enriched in HA-Ago2 IP samples relative to untagged-Ago2 IP or input samples (p ≤ 0.01, fold change ≥ 2). Seventy-five percent of the significantly enriched Ago2 peaks were located in mRNA 3’ UTRs (Figure 2B), consistent with known miRNA activity (Bartel, 2018). A subset of 3’ UTR peaks overlapped seed sites for the 33 active miRNAs from the miR-379/410 cluster. Before investigating miR-379/410 targets, we first analyzed the seed families represented in the cluster. The targeting properties of mature miRNAs are dictated by their seed sequence, a 7-nucleotide sequence that directs RISC to mRNA targets (Bartel, 2018). miRNAs with identical seed sequence are members of a seed family and potentially share mRNA target sites. We grouped the 78 mature miRNAs from the cluster into 70 distinct families based on their 7-mer seed sequences (Figure S2E). We found 60 seed families unique to miR-379/410, and 10 families that share a seed sequence with a miRNA outside the cluster. In the latter category, seed families with multiple expressed family members were excluded from subsequent analysis since repressive activity cannot be conclusively attributed to miR-379/410.

We independently analyzed evolutionarily conserved and nonconserved seed sites with significantly enriched Ago2 peaks, and found that four miRNAs (miR-543-3p, miR-369-3p, miR-154-5p, and miR-495-3p) had the highest number of Ago2-bound conserved seed sites (Figure S2F). Similarly, in a metagene analysis for Ago2 abundance at miRNA seed sites, we observed an enrichment for Ago2 at conserved seed sites for these four miRNAs, but not at nonconserved seed sites (Figure 2C). A similar result was observed in metagene profiles for all Ago2-bound miR-379/410 miRNAs combined (Figure S2G). Using Ago2 seCLIP-seq, we identified miRNAs from the maternally-expressed miR-379/410 cluster that are active in iNs and guide Ago2 to target sites on mRNAs.

### miR-379/410 directs Ago2 to paternally-expressed genes and non-imprinted metabolic regulators

Next, we examined the identity and functional classification of miR-379/410 targets. Given the known functional antagonism between imprinted protein-coding genes, we hypothesized that maternally-expressed miRNAs may direct the RISC complex to paternally-expressed genes. We analyzed metagene profiles of Ago2 abundance at miR-379/410 seed sites on known imprinted genes (Figure 2D). Ago2 was enriched at miR-379/410 seed sites on paternally-expressed genes, but not on maternally-expressed genes. In total, seven paternally-expressed genes are targeted by miR-379/410 (p = 7 × 10^−5^, hypergeometric test), some of which are co-targeted by multiple miRNAs from the cluster (Figure 2E, Figure S2H). *Plagl1*, *Peg3,* and *Mkrn3* are known drivers of paternal programs in the brain that regulate feeding behavior, energy homeostasis, growth, and puberty (Abreu et al., 2013; Curley et al., 2005; Tanaka et al., 2019; Varrault et al., 2006). *Igf2* is a paternally-expressed gene known to control embryonic growth (DeChiara et al., 1991; Ferguson-Smith et al., 1991) but whose function in the postnatal brain remains poorly defined. In fact, several reports indicate *Igf2* switches to maternal-biased expression in the brain (Andergassen et al., 2017; Bonthuis et al., 2015; Perez et al., 2015). Thus, miR-379/410 targeting of *Igf2* may have arisen from regulation in peripheral tissues during early development.

We then looked beyond imprinted targets and analyzed all mRNAs with significant Ago2 binding at miR-379/140 seed sites (Table S1). Ontology analysis of these additional 219 targets indicated they are enriched for functions in diverse metabolic processes, mostly through their roles as transcriptional regulators (Figure 2F). For example, the miR-379/410 targets *Txnip* and *Tbx3* are expressed in energy-sensing neurons and are involved in nutrient regulation (Blouet and Schwartz, 2011; Quarta et al., 2019). Several additional miR-379/410 targets are involved in glucose-responsive pathways, including *Pou2f1* (Shakya et al., 2009), *Sik1* (Berdeaux, 2011), *Hdac4* (Wang et al., 2011b), and *E2f1* (Denechaud et al., 2017). Overall, these data suggest maternally-expressed miR-379/410 acts to repress paternal programs in iNs by recruiting the RISC complex to paternally-expressed genes and additional non-imprinted metabolic regulators.

### Paternally-expressed genes are de-repressed upon maternal miR-379/410 deletion

To determine if miR-379/410 targets are de-repressed upon loss of miR-379/410, we generated cells with maternal-specific deletion of the entire 50 kb miRNA cluster. We co-transfected hybrid ESCs with dual single guide RNAs complementary to sites flanking the miR-379/410 host gene (“*Mirg*”) and loxP-containing donor oligonucleotides. We identified a Mirg^fl/+^ clonal cell line with dual loxP insertion on the maternal allele by Sanger sequencing across allele-specific SNPs, and subsequently deleted the intervening miRNA cluster by transient transfection of Cre recombinase (Figure 3A and Figure S3A). Expression of *Mirg* was completely lost in Mirg^Δ/+^ cells, and no mature miRNAs from the miR-379/410 cluster were produced (Figure 3B, Figure S3B-C). Mirg^Δ/+^ cells exhibited no change in differentiation rate or neuronal morphology (Figure S3D-E). In Mirg^Δ/+^ iNs, we observed loss of miR-379/410 activity by significant de-repression of luciferase reporters containing perfect miRNA target sites (Figure 3C).

**Figure 3.**
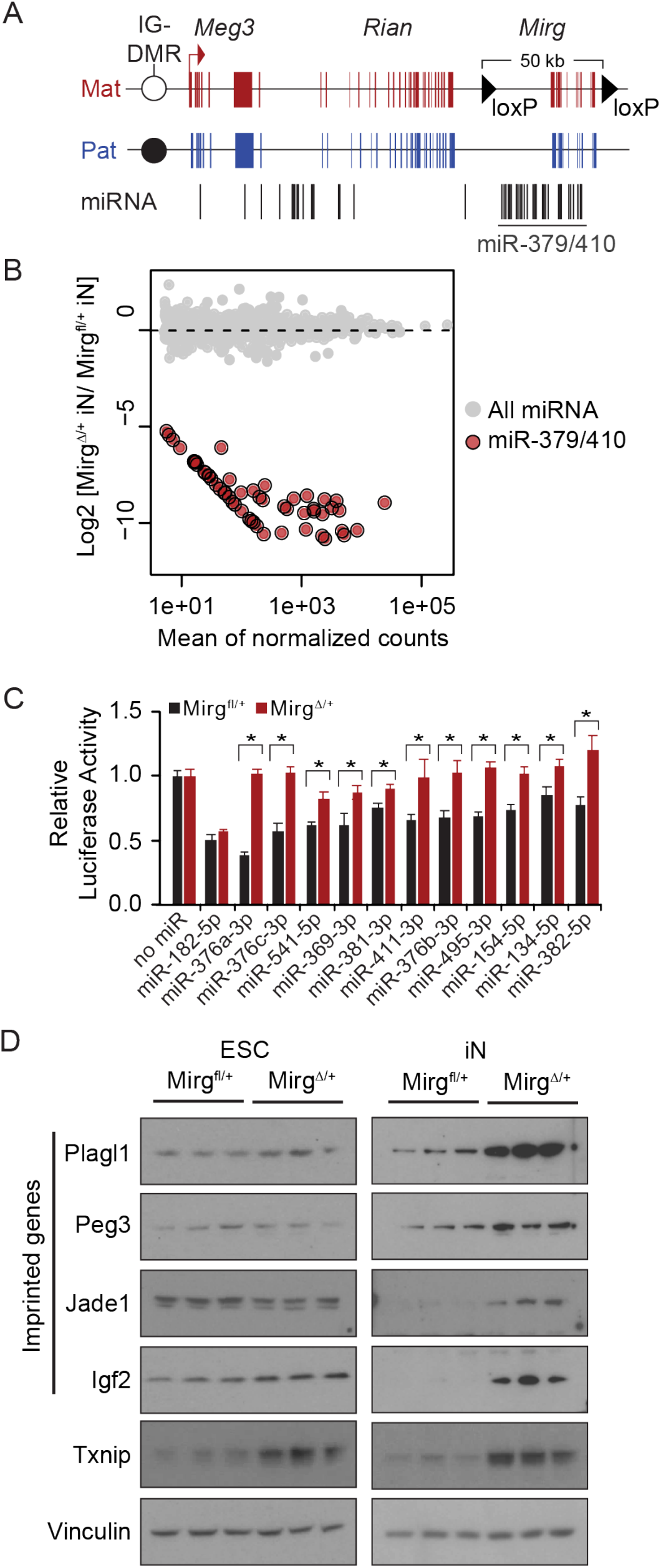
Paternally-expressed genes are de-repressed upon maternal miR-379/410 deletion. **A.** Schematic of the *Meg3* genomic locus showing location of maternal loxP insertions flanking the miR-379/410 cluster. IG-DMR, intergenic differentially methylated region. **B.** Expression changes of miRNAs upon maternal deletion of miR-379/410 shown as an MA plot. Gray, all miRNAs; Red, miRNA-379/410. **C.** Bar graph of Firefly luciferase activity with or without 3’ UTR miRNA target sites relative to Renilla luciferase internal control. Black, Mirg^fl/+^ iNs; Red, Mirg^Δ/+^ iNs; Mean ± SEM (n = 6); *p ≤ 0.05, calculated using unpaired two-tailed Student’s t-test. **D.** Western blot of protein lysates from Mirg^fl/+^ and Mirg^Δ/+^ ESCs and iNs probed with antibodies for Plagl1, Peg3, Jade1, Igf2, and Txnip. Vinculin was used as a loading control. Biological triplicates are shown for each condition. See also Figure S3.

We next analyzed the protein level of miR-379/410 targets identified by Ago2 seCLIP-seq in Mirg^fl/+^ and Mirg^Δ/+^ cells. We observed a neuron-specific up-regulation of the paternally-expressed genes, *Plagl1*, *Peg3*, and *Jade1*, upon maternal miR-379/410 deletion (Figure 3D). The imprinted gene, *Igf2*, also showed robust up-regulation in Mirg^Δ/+^ iNs as well as a slight increase in Mirg^Δ/+^ ESCs. Lastly, the non-imprinted miR-379/410 target *Txnip* that functions in nutrient sensing, was increased in both ESCs and iNs upon maternal miR-379/410 deletion. The stronger de-repression of miR-379/410 targets in iNs compared to ESCs is consistent with our observation of miR-379/410 induction upon differentiation (see Figure 1F). Because imprinted genes are dysregulated in Mirg^Δ/+^ cells, we asked if imprinting at the *Meg3*/*Dlk1* locus is altered by maternal miR-379/410 deletion. A strong paternal bias in methylation at the *Meg3*/*Dlk1* intergenic differentially methylated region (IG-DMR) was observed in both Mirg^fl/+^ and Mirg^Δ/+^ cells (Figure S3F). Accordingly, proper imprinted expression of *Meg3* and *Dlk1* was maintained upon miR-379/410 deletion (Figure S3G), suggesting these miRNAs do not have a cis-regulatory role in *Meg3*/*Dlk1* imprinting.

### miR-379/410 regulates genes involved in feeding behavior and synaptic function

Because miR-379/410 targets are de-repressed in Mirg^Δ/+^ iNs, including transcriptional regulators like *Plagl1*, we analyzed the global transcriptome upon maternal miR-379/410 deletion by rRNA-depleted RNA-seq. We observed 332 differentially expressed genes in Mirg^Δ/+^ iNs (padj ≤ 0.05, fold change ≥ 2) (Figure 4A). We performed gene set enrichment analysis to identify gene ontology terms associated with up-or down-regulated genes. Consistent with the idea that maternally-expressed miRNAs repress a paternal gene program competing for nutrient control, we found that miR-379/410 deletion resulted in up-regulation of genes involved in feeding behavior (Figure 4B-C, top). Mirg^Δ/+^ iNs expressed higher levels of *Mrap2*, *Mchr1*, *Glp1r*, *Npy2r*, and *Galr2*, neuronal genes known to facilitate the effects of nutrient signals on appetite control (Baggio and Drucker, 2014; Georgescu et al., 2005; Naveilhan et al., 1999; Srisai et al., 2017). These data indicate a nutrient responsive pathway is negatively regulated by maternally-expressed miRNAs.

**Figure 4.**
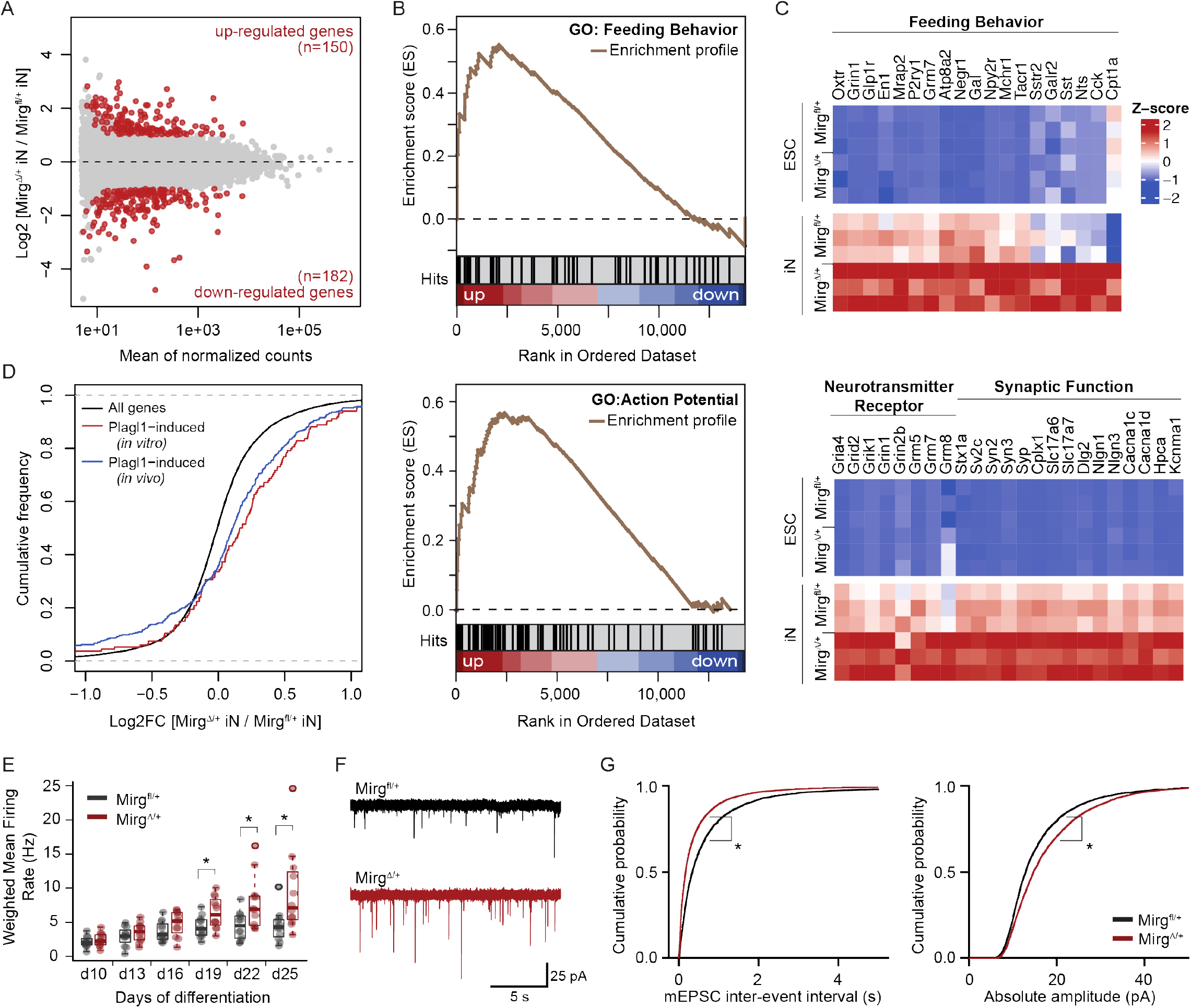
miR-379/410 controls gene programs in feeding behavior and synaptic transmission. **A.** mRNA expression changes in Mirg^Δ/+^ iNs relative to Mirg^fl/+^ iNs shown as an MA plot. Gray, all mRNAs; Red, differentially expressed mRNAs (p ≤ 0.05, fold change ≥ 2). **B.** Enrichment profiles from gene set enrichment analysis for gene ontology (GO) terms “feeding behavior” (normalized enrichment score = 1.76, FDR q-value = 0.18) and “action potential” (normalized enrichment score = 1.88, FDR q-value = 0.11). Genes were ranked from most up-regulated to most down-regulated in Mirg^Δ/+^ iNs relative to Mirg^fl/+^ iNs. **C.** Heatmap of core enriched genes associated with feeding behavior (top) and synaptic activity (bottom). Z-scores were calculated based on expression of each gene in Mirg^fl/+^ and Mirg^Δ/+^ ESCs and iNs (n = 3 for each condition). **D.** Cumulative distribution function (CDF) plot of gene expression upon miR-379/410 deletion for all genes (black, n = 14,325), genes induced by *Plagl1* overexpression in N2A cells (Varrault et al., 2017) (red, n = 134, p = 1.8 × 10^−8^), and genes induced by *Plagl1* overexpression in mouse cortical neurons (Rraklli et al., 2016) (blue, n = 356, p = 8.1 × 10^−11^). P-values were calculated using two-sided Kolmogorov-Smirnov test. **E.**Boxplot of weighted mean firing rate from MEA recordings at day 10 to 25 of differentiation. Individual data points are the mean of 16 electrodes in a single well. Black, Mirg^fl/+^ iNs (n = 12 wells); Red, Mirg^Δ/+^ iNs (n = 12 wells). **F**. Example voltage-clamp recordings of miniature excitatory post-synaptic events (mEPSCs) in Mirg^fl/+^ and Mirg^Δ/+^ iNs. **G**. CDF plot of mEPSC inter-event interval (p = 10^−98^) and amplitude (p = 10^−37^) for n = 4459 and 7752 events in Mirg^fl/+^ and Mirg^Δ/+^ iNs respectively. P-values were calculated using two-sided Kolmogorov-Smirnov test. See also Figure S4.

Additional gene sets associated with neuronal function, such as action potential, membrane depolarization, and fear response, were also enriched in up-regulated genes (Figure 4B, bottom; Figure S4A). Leading edge analysis highlighted a group of neurotransmitter receptors and synaptic proteins up-regulated upon maternal miR-379/410 deletion (Figure 4C, bottom). No gene sets associated with neurological processes were enriched in down-regulated genes. Gene sets enriched for down-regulated genes were driven by a large set of ribosomal protein-coding genes that are subtly down-regulated (fold change = 1.1-1.4) in Mirg^Δ/+^ iNs (Figure S4A).

We next asked if any transcriptional changes in Mirg^Δ/+^ iNs can be attributed to the de-repression of paternally-expressed genes. For example, we identified *Plagl1* as a miR-379/410 target, and *Plagl1* is known to stimulate neuronal gene expression *in vitro* and *in vivo* (Rraklli et al., 2016; Varrault et al., 2017). We analyzed these *Plagl1*-induced genes and found they were significantly up-regulated in Mirg^Δ/+^ iNs (p < 10^−8^) (Figure 4D), consistent with the increased level of Plagl1 protein upon miR-379/410 deletion. Plagl1-induced genes that are up-regulated in Mirg^Δ/+^ iNs include genes associated with synaptic transmission (*Grid2*, *Grik3*, *Pde4a*, *Cacng2*, *Syt7*) and synapse organization (*Synpo*, *Shank1*, *Elfn1*, *Sncb*). Interestingly, the receptor for Glucagon-like peptide-1 (*Glp1r*) involved in neuronal control of feeding behavior (Turton et al., 1996) was up-regulated by *Plagl1* overexpression *in vivo* (Rraklli et al., 2016) and miR-379/410 deletion in iNs (padj ≤ 0.01, fold change = 1.4). Together, these data show that maternally-expressed miRNAs have both primary and secondary effects on nutrient-regulatory pathways.

### Increased frequency and amplitude of synaptic events upon maternal miR-379/410 deletion

Genes involved in synaptic transmission are up-regulated in Mirg^Δ/+^ iNs, so we asked if maternal miR-379/410 deletion affects synaptic activity. Mirg^fl/+^ and Mirg^Δ/+^ iNs were co-cultured with astrocytes under physiological glucose levels and electrophysiology was assessed using two independent techniques, multielectrode array (MEA) and patch clamp. First, electrical activity was measured in Mirg^fl/+^ and Mirg^Δ/+^ iNs by changes in the extracellular field potential on MEA plates (Figure S4B). Mature Mirg^Δ/+^ iNs (day 19 to 25) showed increased electrical activity as indicated by the significant increase in mean firing rate and electrode bursts (Figure 4E, Figure S4C). Second, we performed whole-cell patch clamp recordings on Mirg^fl/+^ and Mirg^Δ/+^ iNs (day 19 to 22). We recorded miniature excitatory post-synaptic activity (mEPSC) by performing voltage-clamp recordings in the presence of tetrodotoxin and bicuculline to block Na^+^ currents and GABA_A_ transmission, respectively (Figure 4F, Figure S4D). In Mirg^Δ/+^ iNs, we observed a decrease in the inter-event interval of mEPSCs, and consequently, an increase in mEPSC frequency (Figure 4G, Figure 4SE). We also observed increased mEPSC amplitude in Mirg^Δ/+^ iNs (Figure 4G, Figure 4SE). Maternal miR-379/410 deletion does not disturb the membrane properties of iNs, as assessed by unaltered Na^+^ and K^+^ conductance, spiking activity, and resting membrane potential (Figure S4F-I). According to both the MEA and patch clamp recordings, these results collectively demonstrated increased excitatory synaptic transmission upon maternal miR-379/410 deletion.

## Discussion

In this study, we investigated the neuronal activity of imprinted miRNAs to better understand why they acquired parental-specific expression. We generated a robust model system for studying imprinted miRNA activity in neurons by Ngn2-induced differentiation of hybrid ESCs. We discovered that maternally-expressed miRNAs from the miR-379/410 cluster repress paternally-expressed genes, including *Plagl1* and *Peg3*, as well as additional metabolic regulators like *Txnip*. Maternal deletion of miR-379/410 de-repressed these targets, resulting in up-regulation of gene programs associated with feeding behavior and synaptic transmission. Lastly, we showed that these changes ultimately lead to increased synaptic signaling. These data establish that maternally-expressed miRNAs increase matrilineal inclusive fitness by antagonizing paternally-driven gene programs in neurons, and lead us to propose that imprinted non-coding RNAs participate in parental genomic conflict.

There are additional single copy and clustered miRNAs transcribed from imprinted domains, but further study is needed to determine the selective forces by which they arose. It also remains to be determined what role, if any, maternal-biased Ago2 expression plays in imprinted miRNA activity. A small group of miRNAs from the upstream region of *Meg3* (miR-127, miR-136, miR-431, miR-433, and miR-434) are transcribed antisense to, and therefore are fully complementary to, the paternally-expressed gene *Rtl1*. There is evidence these miRNAs repress *Rtl1* through the RISC pathway, and this regulation is likely involved in the callipyge phenotype in sheep (Davis et al., 2005). Our findings demonstrate that a large miRNA cluster from the downstream region of *Meg3* repress paternal genes transcribed from distant imprinted loci. The extent to which other imprinted miRNAs regulate imprinted genes ‘in trans’ will be of great interest.

In our study, we found that a subset of the transcriptional changes observed in Mirg^Δ/+^ iNs, including several genes involved in synaptic transmission, can be attributed to overexpression of the paternal transcriptional regulator *Plagl1*. Interestingly, *Plagl1* belongs to a network of co-varying imprinted genes. *Igf2*, a direct transcriptional target of *Plagl1*, *Peg3*, and *Meg3* also belong to this imprinted gene network (Varrault et al., 2006, 2017). Our data shows that miRNAs co-transcribed from the maternal *Meg3* locus repress *Plagl1*, *Peg3*, and *Igf2*, suggesting miR-379/410 may function as a master coordinator of this imprinted network.

It was previously shown that miR-379/410 knockout mice have an impaired response to changes in nutrient availability during the fetal-neonatal transition (Labialle et al., 2014). Defective hepatic gene expression likely accounts, at least in part, for the knockout phenotype, but our data suggests that loss of miRNA activity in the brain could also contribute to the observed impairment in energy homeostasis. Changes in the nutrient environment were previously shown to increase the frequency of action potentials in hypothalamic neurons through an AMPAR-dependent mechanism (Yang et al., 2011). In our patch clamp studies, we recorded mostly AMPA_R_-mediated mEPSCs by blocking NMDA_R_ activity via the addition of 1 mM Mg^2+^ to the bath solution and holding the cells at a negative potential (−70 mV). We observed increased frequency and amplitude of synaptic events in miR-379/410 deletion iNs. While hypothalamic synaptic activity has not been reported in miR-379/410 knockout mice, *in vitro* recordings of hippocampal neurons also showed increased excitatory synaptic transmission (Lackinger et al., 2018). In the future, conditional deletion of maternal miR-379/410 in a stage-specific or neuron subtype-specific manner may help reveal the relative contributions of miRNA dysregulation to metabolic and synaptic phenotypes, and reveal how competition between parental genomes shapes organismal development.

## Acknowledgements

We thank R. Jaenisch for sharing the hybrid ESC line and access to the multielectrode array and pyrosequencer. We thank the Koch Institute’s Robert A. Swanson Biotechnology Center for technical support, specifically the Flow Cytometry Core, the Genomics Core, and the Barbara K. Ostrom Bioinformatics and Computing Facility. This research was supported by NIH grants P01-CA042063 (P.A.S.), R01-GM034277 (P.A.S.), R01-CA133404 (P.A.S.), R01-EY007023 (M.S.), R01-EY028219 (M.S.), R01-NS090473 (M.S.), and the NCI Koch Institute Cancer Center Support (Core) Grant P30-CA14051. Additional support was provided by postdoctoral fellowships NIH F32HD090833 (A.J.W.), FRQS 31677 (V.B.-P.), and NSERC PDF-48724-2016 (V.B.-P.).

## Author Contributions

Conceptualization, A.J.W. and P.A.S.; Methodology, A.J.W. and V.B.-P.; Investigation: A.J.W., H.N.J., and V.B.-P.; Writing-Original Draft, A.J.W., and V.B.-P.; Visualization, A.J.W.; Supervision, M.S. and P.A.S.; Funding Acquisition, A.J.W., V.B.-P., M.S., and P.A.S.

## STAR Methods

### CONTACT FOR REAGENT AND RESOURCE SHARING

Further information and requests for resources and reagents should be directed to and will be fulfilled by the Lead Contact, Phillip A. Sharp.

### EXPERIMENTAL MODEL AND SUBJECT DETAILS

#### Cell culture

##### Cell line maintenance

The ESC line used in this study was derived from an F1 *Mus. musculus* × *Mus. castaneous* cross (F_1.2-3_, (Eggan et al., 2001)). ESCs were adapted to feeder-free conditions by 10 passages on tissue culture plates pre-coated with 0.2% gelatin in phosphate-buffered saline (PBS). For routine passaging, cells were washed in HEPES buffered saline (HBS), dissociated using 0.25% trypsin-EDTA, then plated on gelatin-coated plates in ESC medium [Dulbecco’s Modified Essential Media (Thermo) containing 10 mM HEPES (Thermo), 15% HyClone fetal bovine serum (Thermo), 1000 U/mL leukemia inhibitory factor (Millipore Sigma), 0.11 mM ß-mercaptoethanol (Sigma), 1X non-essential amino acids (Thermo), 2 mM L-glutamine (Thermo), 1X penicillin streptomycin solution (Corning)]. Cells were maintained in a humidified 5% CO_2_ incubator at 37°C.

##### Neuron differentiation

ESC lines stably expressing rtTA and TetO-Ngn2 (see below) were cultured for 24 hours on gelatin-coated tissue culture plates in N2B27 medium [1:1 ratio of DMEM/F12 + Glutamax and Neurobasal Medium (Thermo), supplemented with 1X N2 supplement (Thermo), 1X B27 supplement minus Vitamin A (Thermo), 0.5X non-essential amino acids (Thermo), 1 mM L-glutamine (Thermo), 1X penicillin streptomycin solution (Corning), 0.05 mM ß-mercaptoethanol (Sigma), 0.5 mM sodium pyruvate (Thermo), 0.02% BSA Fraction V (Thermo), 0.25 ug/mL laminin (Thermo), 10 ng/mL BDNF (Sigma), 10 ng/mL NT3 (R&D Systems), 1 ug/mL doxycycline (Sigma)]. Then, cells were dissociated using accutase (STEMCELL Technologies) and 6 × 10^5^ cells were plated on PEI-coated six-well plates (0.1% PEI in borate buffer). Borate buffer contains 50 mM boric acid (Sigma) and 24 mM sodium tetraborate (Sigma) at pH 8.4. One half of the medium was replaced every two days. iNs were harvested at day 10 unless otherwise noted.

### METHODS

#### Generation of EF1α-M2rtTA;TetO-Ngn2-mCherry;Tubb3-GFP cell line

First, a gBlock gene fragment (IDT) was synthesized of codon-optimized P2A-eGFP-T2A-Puro sequence flanked by 450 bp homology arms surrounding the *Tubb3* stop codon and cloned into pUC19 plasmid (Addgene). A single guide RNA (sgRNA) targeted to the Tubb3 3’ UTR (Tubb3sg Fwd, 5’-CACCGACCAGCTCATTAGGGCTCCC-3’; Tubb3sg Rev, 5’-AAACGGGAGCCCTAATGAGCTGGTC-3’) was cloned into pX458 (Addgene), and co-transfected with the homology directed repair template into F_1.2-3_ ESCs using Lipofectamine 2000 (Thermo). Transfected cells were re-plated at a low density and single cell clones were manually isolated. Genomic DNA was prepared using Quick Extract (Epicentre) and a single cell clone with successful insertion of Tubb3-P2A-GFP-T2A-Puro was confirmed by PCR. Second, the piggybac pAC4 vector (co-expressing EF1α-M2rtTA and CMV-Hygro) was co-transfected with mPBase (expressing the PiggyBac transposase) into Tubb3-GFP ESCs. Cells with stable integration of EF1α-M2rtTA were selected in the presence of 150 ug/mL Hygromycin B for 10 days. Third, the lentiviral pTetO-Ngn2-Puro plasmid (Addgene) was modified to pTetO-Ngn2-mCherry using Gibson Assembly (NEB). Lentivirus was produced in HEK293 cells by co-transfection of pTetO-Ngn2-mCherry, psPAX2(Δ8.2), and pCMV-VSV-G using TransIT-LT1 Transfection Reagent (Mirus). Viral-containing supernatant was applied to EF1α-M2rtTA;Tubb3-GFP ESCs in the presence of 5 ug/mL polybrene (Sigma). An infected single cell clone (EF1α-M2rtTA;TetO-Ngn2-mCherry;Tubb3-GFP) was identified and confirmed by mCherry/GFP fluorescence following doxycycline treatment.

#### Generation of Mirg^fl/+^ and Mirg^Δ/+^ cell lines

sgRNAs (Mirg1sg and Mirg2sg) were designed to sites flanking the miR-379/410 cluster and were cloned into the pX458 plasmid (Addgene). sgRNA sequences are as follows: Mirg1sg Fwd 5’-CACCGTCTATCGGGAGGATAGTACA-3’; Mirg1sg Rev 5’-AAACTGTACTATCCTCCCGATAGAC-3’; Mirg2sg Fwd, 5’-CACCGATAGTCTGATTCAAGTGCCG-3’; Mirg2sg Rev, 5’-AAACCGGCACTTGAATCAGACTATC-3’. Templates for homology directed repair (HDR) each contain a loxP site flanked by 60 bp homology arms. sgRNAs and HDR templates were co-transfected into EF1α-M2rtTA;TetO-Ngn2-mCherry;Tubb3-GFP ESCs using Lipofectamine2000 (Thermo). Single cells were isolated by FACS into individual wells of a 96-well plate and screened for loxP insertion by PCR at the upstream and downstream insertion sites. A single cell clone with dual loxP insertion was confirmed to be maternal-specific by Sanger sequencing of allelic SNPs within loxP-containing PCR products. Next, the miR-379/410 cluster was deleted by transient transfection of pCAG-Cre:GFP (Addgene). Since cell clones were isolated by FACS for GFP-negative (Mirg^fl/+^) and GFP-positive (Mirg^Δ/+^) cells, and Cre-mediated deletion of miR-379/410 was confirmed by PCR.

#### Generation of untagged-Ago2 and HA-Ago2 cell lines

Untagged-Ago2 or HA-Ago2 was cloned into the multiple cloning site of PB533A-2 (System Biosciences) by restriction enzyme ligation, and subsequently co-transfected with mPBase (expressing the PiggyBac transposase) into Mirg^fl/+^ ESCs. Cells with stable Ago2 integration were selected in neomycin-containing medium for 10 days.

#### Flow cytometry and fluorescence activated cell sorting (FACS)

Single cells were sorted using the FACSAria cell sorter (Becton Dickinson) into individual wells of a gelatin-coated 96-well plate containing ESC medium. To analyze the efficiency of neuron differentiation, cells were analyzed on the FACSCelesta (Becton Dickinson) flow cytometry instrument with FACSDiva v8.0 software. FCS files were exported and analyzed with FlowJo v9.

#### Fluorescence Imaging

Cells were imaged on the Incucyte (Sartorius) live cell imager or ApoTome.2 (Zeiss) fluorescence microscope using the brightfield and mCherry channels.

#### Luciferase assay

Perfect target sites fully complementary to mature miRNAs were cloned into the multiple cloning site in pmirGLO dual luciferase expression vector (Promega) using restriction enzyme ligation and plasmid insertions were verified by sequencing. Mirg^fl/+^ and Mirg^Δ/+^ ESCs were plated on gelatin-coated, white-walled 96-well plates at a density of 8,000 cells per well. After 24 hours, cells were transfected with 200 ng pmirGLO plasmid using Lipofectamine2000. Transfection medium was replaced after six hours with N2B27 medium containing 1 ug/mL doxycycline to induce differentiation. After 48 hours of differentiation, the luciferase assay was performed using the reagents and protocol from the Dual-Glo Luciferase Assay System (Promega) according to the manufacturer’s instructions. Luminescence was measured on the Infinite 200 (Tecan Life Sciences). For each sample, Firefly luminescence was normalized to Renilla luminescence as an internal expression control. The Firefly/Renilla ratios for each sample were then normalized to pmirGLO containing no miRNA binding site. Six biological replicates were included for each condition.

#### miRNA qRT-PCR

Total RNA was harvested using TRIzol (Thermo) according to the manufacturer’s instructions. Reverse transcription was performed using the miScript II RT Kit (Qiagen) with 500 ng RNA and the miScript HiSpec Buffer, followed by PCR using the miScript SYBR Green PCR Kit (Qiagen). miRNAs were normalized relative to U6. The mean and standard error was calculated from biological triplicates. miRNA-specific forward primers were the mature miRNA sequence, as follows: miR-409-3p, 5’-GAATGTTGCTCGGTGAACCCCT-3’; miR-376b-3p, 5’-ATCATAGAGGAACATCCACTT-3’; miR-541-5p, 5’-AAGGGATTCTGATGTTGGTCACACT-3’; miR-134-5p, 5’-TGTGACTGGTTGACCAGAGGGG-3’; miR-381-3p, 5’-TATACAAGGGCAAGCTCTCTGT-3’; miR-379-5p, 5’-TGGTAGACTATGGAACGTAGG-3’; miR-300-3p, 5’-TATGCAAGGGCAAGCTCTCTTC-3’; miR-411-5p, 5’-TAGTAGACCGTATAGCGTACG-3’; miR-154-5p, 5’-TAGGTTATCCGTGTTGCCTTCG-3’; miR-382-5p, 5’-GAAGTTGTTCGTGGTGGATTCG-3’; miR-369-5p, 5’-AGATCGACCGTGTTATATTCGC-3’; miR-369-3p, 5’-AATAATACATGGTTGATCTTT-3’; miR-770-3p, 5’-CGTGGGCCTGACGTGGAGCTGG-3’; U6, 5’-CGCTTCGGCAGCACATATAC-3’.

#### Northern blot

Total RNA was harvested using TRIzol (Thermo) according to the manufacturer’s instructions. Four micrograms total RNA was diluted in Gel Loading Buffer II (Thermo), heat denatured at 95°C for 5 minutes, separated on an 8% urea gel (National Diagnostics), and then transferred by semi-dry transfer (300 mA, 60 min) to Amersham Hybond N+ membrane (GE Healthcare). The Northern probes were 5’-end labelled with ATP Gamma ^32^P (Perkin Elmer) using T4 polynucleotide kinase (New England Biolabs), and then hybridized to the membrane overnight in ULTRAhyb-Oligo buffer (Thermo) in a 42°C rotating hybridization oven. The membrane was washed in 2X SSC containing 0.1% SDS three times and then exposed to a PhosphorImager. The oligonucleotide probe sequences were miR-411-5p, 5’-CGTACGCTATACGGTCTACTA-3’; miR-410-3p, 5’-ACAGGCCATCTGTGTTATATT-3’; miR-376a-3p, 5’-CGTGGATTTTCCTCTACGAT-3’; 5.S rRNA, 5’-TCCTGCAATTCACATTAATTCTCGCAGCTAGC-3’.

#### Western blot

Whole cell extract was prepared by washing Mirg^fl/+^ and Mirg^Δ/+^ cells in ice cold 1X phosphate buffered saline (PBS), then lysing in pre-chilled RIPA buffer containing 10 mM Tris-HCl pH 7.4, 100 mM NaCl, 1% Trixon X-100, 0.1% SDS, 1 mM EDTA, and cOmplete, EDTA-free protease inhibitor cocktail (Roche). Cell lysates were rotated for 30 minutes at 4°C, then cell debris was pelleted at 4°C for 30 minutes at 12,000 rpm. Protein concentration was determined using the BCA assay (Pierce). 30 ug protein lysate were diluted in 1X NuPAGE LDS Sample Buffer (Thermo) and 1X NuPAGE Reducing Reagent (Thermo), heat denatured at 70 °C for 10 minutes, and separated on NuPAGE 4-12% Bis-Tris gels in 1X NuPAGE MOPS SDS Running Buffer (Thermo). Following transfer to PVDF membrane in 1X NuPAGE Transfer Buffer (Thermo), membranes were blocked in 5% milk in PBST, and incubated overnight at 4C in primary antibody [anti-HA high affinity antibody (Roch 11867423001), Enolase I (CST 3810S), Vinculin (Sigma V9131), anti-Plagl1 (Abcam, ab90472), anti-Peg3 (Abcam, ab99252), anti-Jade1 (Novus, NBP1-83085), anti-Igf2 (Thermo, PA5-71494), anti-Txnip (Cell Signaling Technology, 14715S), anti-Ago2 (Cell Signaling Technology, 2897S)]. Membranes were washed in 1X PBST, then incubated at room temperature for 1 hour in HRP-conjugated secondary antibody [anti-mouse IgG (GE Healthcare NA931V), anti-rabbit IgG (GE Healthcare NA934V)]. Blots were exposed with Western Lightning Plus-ECL (Perkin Elmer) or SuperSignal West Dura Extended Duration Substrate (Thermo).

#### RIP-qRT-PCR

Mirg^fl/+^;untag-Ago2 and Mirg^fl/+^;HA-Ago2 cells were plates on PEI-coated plates (3 × 10^5^ cells per 10 cm plate, see neuron differentiation protocol) and differentiated for six days. Cells were washed in ice cold 1X PBS and lysed for 10 minutes in ice cold lysis buffer [50 mM Tris-HCl pH 7.4, 100 mM NaCl, 1% NP-40, 0.1% SDS, 0.5% sodium deoxycholate, 0.4 U/uL murine RNAse inhibitor (NEB), 1:200 Protease Inhibitor Cocktail (Millipore)]. Cellular debris was pelleted by centrifugation at 14,000 rpm for 10 minutes at 4°C. 50 ul of anti-HA magnetic beads (Thermo) were blocked in 3% BSA and then incubated with cell lysate at 4°C overnight. Beads were separated with a magnet, washed three times in 500 ul cold lysis buffer, and then resuspended in TRIzol. Following RNA isolation, miRNA immunoprecipitation was measured by qRT-PCR (see miRNA qRT-PCR protocol). Fold enrichment in HA-Ago2 iNs was calculated relative to untag-Ago2 iNs.

#### Allele-specific DNA methylation

Genomic DNA was isolated from Mirg^fl/+^ and Mirg^Δ/+^ cells using the GenElute Mammalian Genomic DNA Miniprep Kit (Sigma) according to the manufacturer’s instructions. Bisulfite conversion was performed on 500 ng genomic DNA using the EpiTect Bisulfite Kit (Qiagen). M.SssI-treated DNA was used as a positive control for complete bisulfite conversion. PCR was performed on 20 ng bisulfite converted DNA with primers in the IG-DMR that amplify seven bi-allelic CpG sites and one paternal-specific CpG sites (Fwd, 5’-ATTGGGTTTTGAGGAGTAGTATATTA-3’; 5’ biotinylated Rev, 5’- AAAACACCAAATCAAAACTATTATATCAC-3’) using the PyroMark PCR Kit (Qiagen). Sequencing assays were run on the PyroMark Q48 pyrosequencer (Qiagen) using the sequencing primer 5’-ATTATAGTTATAGATAATAAAGTA-3’. The percent methylation at the paternal-specific CpG site was subtracted from the average percent methylation at non-allelic CpG sites to yield the percent methylation on the maternal allele.

#### Allele-specific qRT-PCR

Taqman SNP genotyping assays (Thermo) were designed to allele-specific SNPs in *Meg3* (Fwd, 5’-GAAGAGCAGCTGGCATTGC-3’; Rev 5’-CGCGCTGGGCATCCT-3’; FAM-labelled maternal probe, 5’-CCATGCCGGCTGAAGA-3’; VIC-labelled paternal probe, 5’-CCATGCCGACTGAAGA-3’) and *Dlk1* (Fwd, 5’-TCGGCCACAGCACCTATG-3’; Rev, 5’-CAGCCTCGCAGAATCCATACT-3’; FAM-labelled maternal probe, 5’-ACCCTGTGACCCCC-3’; VIC-labelled paternal probe, 5’-CCTGCGACCCCC-3’). One-step qRT-PCR was performed using 25 ng DNase-digested RNA from Mirg^fl/+^ and Mirg^Δ/+^ cells with EXPRESS One-Step SuperScript qRT-PCR Kit (Thermo) on the StepOne qPCR machine (Applied Biosystems). The Ct values for maternal and paternal amplification were normalized to a standard curve that was generated using known ratio mixtures of maternal-specific and paternal-specific synthetic amplicons (IDT).

#### Multielectrode array (MEA)

Primary mouse astrocytes (ScienCell) were expanded on tissue culture flasks pre-coated with 50 ug/mL poly-L-lysine for 3-4 passages in Astrocyte Medium-animal (ScienCell). Classic MEA 48 plates (Axion Biosystems) were pre-coated with 0.1% PEI, then 7 × 10^5^ astrocytes and 7 × 10^5^ Mirg^fl/+^ or Mirg^Δ/+^ day 1 iNs were mixed and plated in each well by drop placement. For electrophysiology experiments, iNs were cultured in BrainPhys Neuronal medium (StemCell Technologies) containing 10 ng/mL BDNF (Sigma), 1X N2 supplement (Thermo), SM1 Neuronal Supplement (StemCell Technologies), and 1 ug/mL doxycycline (Sigma). Twelve wells were plated per condition. Recordings of spontaneous activity were performed for 10 minutes at 37°C every 3 days from day 10 to 25 on the MEA Maestro system (Axion Biosystems). Electrode bursts were calculated using the ISI threshold algorithm with minimum number of spikes = 5 and maximum ISI = 100 ms.

#### Patch clamp electrophysiology

1 × 10^5^ primary mouse astrocytes and 2 × 10^5^ Mirg^fl/+^ or Mirg^Δ/+^ day 1 iNs were mixed and plated on PEI-coated 15 mm circular coverslips in a 12-well plate. For electrophysiology experiments, iNs were cultured in BrainPhys Neuronal medium (StemCell Technologies) containing 10 ng/mL BDNF (Sigma), 1X N2 supplement (Thermo), SM1 Neuronal Supplement (StemCell Technologies), and 1 ug/mL doxycycline (Sigma). At day 19 to 22, coverslips were placed in a recording chamber (Series 20, Warner Instruments) and perfused at a rate of 1-2 mL/min with room-temperature ACSF containing the following: 140 mM NaCl, 5 mM KCl, 10 mM HEPES, 1 mM MgSO_4_, 2 mM CaCl_2_ and 10 mM D-Glucose; pH 7.3-7.4 and osmolarity 300. We used 3-6 MΩ glass pipettes filled with K-Glu-based intracellular solution containing 140 mM K-gluconate, 10 mM KCl, 10 mM HEPES, 1 mM EGTA, 0.1 mM CaCl_2_, 1 mM Mg-ATP, 1 mM Na_2_-GTP, and 4 mM D-Glucose; pH 7.4 and osmolarity 290. iNs were visualized with differential interference contrast (DIC) or fluorescence using a charge-coupled device (CCD) camera (ORCA-R2 C10600, Hamamatsu). Recordings were performed with a Multiclamp 700b (Molecular Devices) amplifier, digitized at 10 kHz, and low-pass filtered at 1 kHz with a Digidata 1440 and pClamp10 software (Molecular Devices). To isolate mEPSCs, we added 1 μM TTX (ab120055, Abcam) and 20 μM bicuculline (0131, Tocris) to the bath solution to block Na^+^ and GABA_A_ currents respectively and cells were voltage-clamped at a membrane potential of −70 mV. Access resistance was 17.6±0.7 and 17.4±0.9 MΩ in Mirg^fl/+^ and Mirg^Δ/+^ iNs, respectively. mEPSCs, membrane properties, and spiking properties were analyzed using Clampfit software (version 10.7, Molecular Devices) and custom MATLAB scripts (version R2018b, MathWorks). mEPSCs properties were calculated over 3-minute-long recordings. Spontaneous spiking rate and resting membrane potential were calculated over 1-minute-long current-clamp recordings. Evoked spike rate activity was calculated by counting the number of spikes elicited by injecting +50 pA currents during 1 s. Sodium and potassium currents were measured as the peak negative current and the average positive current following the injection of voltage steps (ranging from −90 to 50 mV; 10 mV increments).

#### Neuron differentiation for sequencing libraries

Mirg^fl/+^ or Mirg^Δ/+^ were differentiated according to the neuron differentiation protocol above and total RNA was prepared from three biological replicates of day 10 iNs from each cell line using TRIzol. RNA was treated with DNase I (Thermo) under standard reaction conditions. DNase-treated RNA was used for preparation of small RNA-seq and rRNA-depleted RNA-sequencing libraries.

#### Small RNA-seq, library preparation

Small RNA-seq libraries were prepared from 100 ng DNase-treated RNA using NEB small RNA-sequencing kit (E7300S) using 16 cycles of PCR amplification. Quality control of final libraries was assessed with a fragment analyzer and qPCR for colony forming units prior to pooling and loading on an Illumina FlowCell (HiSeq2000, 40 bp SE reads). Each sample library was sequenced to a depth of 9-12 M reads.

#### Small RNA-seq, data analysis pipeline

Adapter sequences were first trimmed from small RNA-sequencing reads using cutadapt (v1.4.2) with the following command: cutadapt -a AACTGTAGGCACCATCAAT--minimum length 14 $input/ID_sequence.fastq > $input/ID_sequence_trim.fastq 2> $input/TrimReport-ID.txt

Before mapping small RNA-seq reads to the transcriptome, we added all mature miRNA annotations from miRBase (v21) into the mm10 Gencode M15 gtf file. Mature miRNAs were inserted as distinct transcripts of the appropriate parent pri-miRNA gene annotation, allowing STAR alignment and RSEM counting of gene-level and mature miRNA-level counts. STAR (v2.4.1d) was used to map small RNA-sequencing reads to mm10 genome and transcriptome (Gencode M15 assembly containing miRBase annotations) with the following command: STAR --runThreadN 8 --runMode alignReads --genomeDir $genome/ --readFilesIn $input/ID_sequence_trim.fastq --outFileNamePrefix $input/ID --outReadsUnmapped Fastx --outSAMtype BAM SortedByCoordinate --outFilterMismatchNoverLmax 0.05 --outFilterMatchNmin 16 --outFilterScoreMinOverLread 0 --outFilterMatchNminOverLread 0 --alignIntronMax 1 --sjdbGTFfile $gtf/gencode.vM15.miRBase.gtf --sjdbOverhang 74 --quantMode TranscriptomeSAM

RSEM (v1.2.30) was used to count transcriptome mapped reads with the following command: rsem-calculate-expression --bam --forward-prob 0.5 -p 8 --no-bam-output --calc-pme --seed-length 16 $input/IDAligned.toTranscriptome.out.bam $genome/GRCm38_GencodeM15_miRBase $input/ID

Quantification of RSEM counts was performed on ID.isoforms.results files using DESeq2. Only mature miRNAs with mean normalized counts ≥ 5 were included in the analysis. Differentially expressed miRNAs between Mirg^fl/+^ and Mirg^Δ/+^ iNs were determined as padj ≤ 0.05 and fold change ≥ 2. Heatmap of Log2[Normalized counts] were generated using ComplexHeatmap (v1.20.0) package in R. The proportion of total miRNA reads mapping to miR-379/410 miRNAs was calculated in R using the posterior mean counts from RSEM for all mature miRNAs.

#### rRNA-depleted RNA-seq, library preparation

rRNA-depleted RNA-sequencing libraries were prepared from 400 ng DNase-treated RNA using KAPA RNA HyperPrep Kit with RiboErase (HMR) (Kapa Biosystems) using 9 cycles of PCR amplification. Quality control on final libraries was assessed with a fragment analyzer and qPCR for colony forming units prior to pooling and loading on an Illumina FlowCell (NextSeq 500, 75 bp PE reads). Each sample library was sequenced to a depth of 38-50 M reads.

#### rRNA-depleted RNA-seq, data analysis pipeline

STAR (v2.4.1d) was used to map RNA-sequencing reads to mm10 genome and transcriptome (Gencode M15 assembly) with the following command: STAR --runThreadN 8 --runMode alignReads --genomeDir $genome/ --readFilesIn $input/ID_1.fastq $input/ID_2.fastq --outFileNamePrefix $input/ID --outFilterType BySJout --outReadsUnmapped Fastx --outSAMtype BAM SortedByCoordinate –outFilterMultimapNMax 20 --outFilterMismatchNmax 999 --outFilterMismatchNoverLmax 0.04 --alignIntronMin 70 --alignIntronMax 1000000 --alignMatesGapMax 1000000 --alignSJoverhangMin 8 --alignSJBoverhangMin 1 –sjdbGTFfile $genome/gencode.vM15.primary_assembly.annotation.gtf --sjdbOverhang 74 --twopassMode Basic --quantMode TranscriptomeSAM

RSEM (v1.2.30) was used to count transcriptome mapped reads with the following command: rsem-calculate-expression --bam --paired-end --forward-prob 0 -p 8 --no-bam-output --calc-pme $input/IDAligned.toTranscriptome.out.bam $genome/ $input/rsem/ID

Gene level quantification was performed on RSEM ID.genes.results files using DESeq2. Only genes with mean normalized counts ≥ 5 were included in the analysis. Differentially expressed genes between Mirg^fl/+^ and Mirg^Δ/+^ iNs were determined as padj ≤ 0.05 and fold change ≥ 2.Gene set enrichment analysis (GSEA) was performed by ranking all genes from most up-regulated to most down-regulated and enrichment was calculated for the Molecular Signatures Database v6.2 C5 GO gene sets. For the heatmap of GSEA core enriched genes, we first calculated the z-score across samples for each gene based on the transcripts per million (TPM) output from RSEM using the scale function in R. Then, the heatmap was generated using ComplexHeatmap (v.1.20.0) R package. The cumulative frequency plot (Figure 4D) was generated in R (v.3.5.1) using the plot.ecdf function. Plagl1-regulated genes *in vitro* (Varrault et al., 2017) or *in vivo* (Rraklli et al., 2016) were all induced genes from overexpression of Plagl1 with padj ≤ 0.01 and fold change ≥ 2.

#### seCLIP-seq, library preparation

15 × 10^6^ Mirg^fl/+^;untag-Ago2 or Mirg^fl/+^;HA-Ago2 ESCs were plated on PEI-coated 15 cm plates and differentiated in N2B27 medium containing 1 ug/mL doxycycline for 10 days (2 biological replicates per cell line). Cells were washed in 1X PBS and then UV crosslinked at 400 mJ/cm2. Immunoprecipitation was performed with 100 uL anti-HA magnetic beads per sample (Pierce) overnight at 4°C. seCLIP-seq libraries were prepared according to (Van Nostrand et al., 2017) with TruSeq LT adapters. Input and IP samples were amplified with 9 and 18 cycles of PCR, respectively. Quality control on final libraries was assessed with a fragment analyzer and qPCR for colony forming units prior to pooling and loading on an Illumina FlowCell (HiSeq2000, 40 bp SE reads). Each sample library was sequenced to a depth of 20-30 M reads.

#### seCLIP-seq, data analysis pipeline

Adapter sequences were trimmed from seCLIP-sequencing reads using fastxtoolkit (v0.0.13), followed by removal of PCR duplicated based on unique molecular identifier (UMI), and then the UMI was trimmed with the following commands: fastx_clipper -a AGATCGGAAGAGCACACGTCTGAACTCCAGTCAC -Q33 -l 15 -v –I $input/ID_sequence.fastq -o $input/ID_sequence_clip.fastq fastx_collapser -Q33 -v -i $input/ID_sequence_clip.fastq -o $input/ID_sequence_collapse.fastq fastx_trimmer -f 11 -i $input/ID_sequence_collapse.fastq -o $input/ID_sequence_trim.fastq

To determine which miRNAs were enriched for Ago2 binding, STAR (v2.5.3a) was used to map seCLIP-sequencing reads to mm10 genome (Gencode M15 assembly) with the following command (allowing for reads to map up to 5 locations since some miRNAs are multi-copy in the genome): STAR --runThreadN 8 --runMode alignReads --genomeDir $genome/ --readFilesIn $input/ID_sequence_trim.fastq --outFileNamePrefix $input/ID_ --outReadsUnmapped Fastx --outFilterMultimapNmax 5 --outFilterMultimapScoreRange 1 --outSAMtype BAM SortedByCoordinate --outFilterType BySJout --outFilterScoreMin 10 --alignEndsType EndToEnd --sjdbGTFfile $genome/gencode.vM15.primary_assembly.annotation.gtf Biological replicates were merged using samtools. Then for each mature miRNA annotated in miRbase, enrichment for Ago2 binding was calculated in HA-Ago2 IP samples relative to HA-Ago2 Input using seCLIP perl scrips from the Yeo lab (https://github.com/YeoLab/eclip).

To determine which mRNA targets were enriched for Ago2 binding, the same command as above was used by only uniquely mapped reads were considered (by setting --outFilterMultimapNmax = 1). Biological replicates were merged using samtools. Peaks were called on HA-Ago2 IP sample using the CLIPper tool with the following command: clipper -b $path/HAAgo2IP.merged.bam -s mm10 -o $path/HAAgo2.peaks.bed Then, enrichment for Ago2 binding at each peak was calculated in HA-Ago2 IP samples relative to HA-Ago2 Input and untag-Ago2 IP using seCLIP perl scrips from the Yeo lab (https://github.com/YeoLab/eclip).

#### Metagene analysis

Genome (mm10) coordinates for target seed sites of miR-379/410 miRNAs were downloaded from TargetScanMouse (release 7.2). Metagene binding plots were generated from Ago2 seCLIP-seq binding profiles. “.bam” files from merged biological replicates were loaded into the R package metagene (v2.14.0), and coverage (RPM) was calculated at the seed site and the flanking 500 nt. The metagene profile for imprinted genes contains all conserved, predicted seed sites active miR-379/410 in maternally-expressed or paternally-expressed protein-coding genes from (Tucci et al., 2019).

#### Biological network of miR-379/410 targets using Cytoscape

The biological network of gene ontology terms associated with miR-379/410 targets identified from Ago2 seCLIP-seq were analyzed using the Cytoscape (v3.7.1) app, BiNGO. All genes containing significantly enriched Ago2 peaks overlapping a seed site of an active miR-379/410 miRNA were used as the input, and overrepresentation of gene ontology terms was calculated using hypergeometric test and Benjamini & Hochberg FDR correction relative to the reference set of all gene expressed in iNs.

#### Analysis of in vivo Mirg and miR-379/410 expression

microRNA-seq and polyA RNA-seq data from embryonic day 14.5 and postnatal day 0 mouse tissues were downloaded from the ENCODE project (see GSE82864, GSE82558, and GSE78374 for forebrain samples, and related GSE entries for other tissues). Small RNA-seq data from adult mouse tissues were downloaded from GSE21630 (Kuchen et al., 2010). polyA RNA-seq from isolated cortical cell types were downloaded from GSE52564 (Zhang et al., 2014). Raw .fastq files from these publicly available datasets were analyzed according to the small RNA-seq and RNA-seq data analysis pipelines described above.

### QUANTIFICATION AND STATISTICAL ANALYSIS

For all analysis with p values, significance was determined at p ≤ 0.05 and values are shown. Hypergeometric statistical test for enrichment was used to calculate the enrichment of miR-379/410 seCLIP targets for paternally-expressed genes (N = 13,825 expressed protein-coding genes, n = 63 paternally-expressed protein-coding genes, K = 227 miR-379/410 targets from seCLIP-seq, k = 7 miR-379/410 paternally-expressed targets). Box-whisker plots were generated using R with the box extending from the 25^th^ to 75^th^ percentile and a thick bar at the 50^th^ percentile. Dashes lines extend from the min to max data points with outliers shown. For CDF analysis, significance was measured using two-sided Kolmogorov-Smirnov (KS) test using the ks.test function in R.

### DATA AND CODE AVAILABILITY

The rRNA-depleted RNA-seq, small RNA-seq, and seCLIP-seq data are in the process of being deposited in NCBI’s Gene Expression Omnibus and will be available for review.

## Supplemental Tables

**Table S1.** HA-Ago2 seCLIP-seq peaks that overlap an active miR-379/410 seed site and are significantly enriched above untag-Ago2 IP and HA-Ago2 input samples.

## Supplemental Figures and Figure Legends

**Figure S1.**
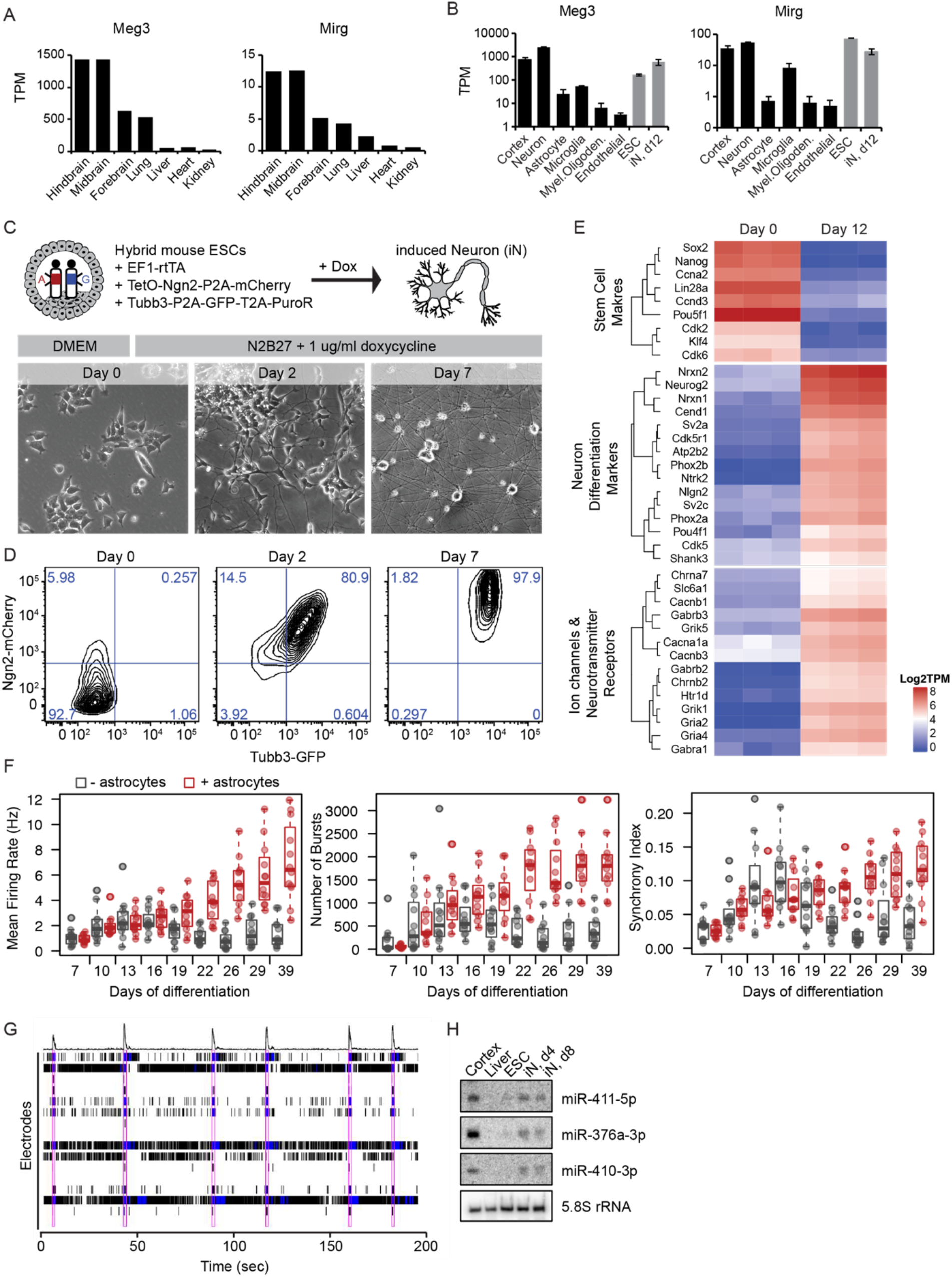
An *in vitro* model system for studying imprinted gene function in neurons. **A.** Quantification of *Meg3* and *Mirg* expression as transcripts per million (TPM) from postnatal day 0 mouse tissues. **B.** TPM quantification of *Meg3* and *Mirg* expression from cell types isolated from the mouse cortex, in vitro ESCs, and iNs (day 12). Mean ± abs error (n = 2) for neuron, astrocyte, microglia, myelinating oligodendrocyte, and endothelial samples; Mean ± std deviation (n = 3) for whole cortex, ESCs, and iNs. **C.** Schematic overview of Ngn2-induced differentiation and brightfield images of ESCs and iNs (day 2 and 7). **D.** Fluorescence activated cells sorting (FACS) of ESCs and iNs (day 2 and 7) based on Ngn2-mCherry and endogenously tagged Tubb3-GFP. Percentage of cells in each gated quadrant is indicated. **E.** Heatmap of transcripts per million (TPM) from rRNA-depleted RNA-seq of ESCs (day 0, n = 3) and iNs (day 12, n = 3) for stem cell markers, neuron differentiation markers, and ion channels or neurotransmitter receptors. **F.** Analysis of multielectrode array (MEA) recordings for mean firing rate (Hz), number of electrode bursts, and synchrony index across a time course of differentiation. Black, iNs alone; Red, iNs co-cultured with astrocytes. Individual data points are the mean of 16 electrodes in an individual well, and boxplot summarize the data of 12 wells per condition. **G.** An example raster plot of iNs (day 19). The first row is the population histogram and all subsequent rows are tracks of the 16 individual electrodes in the well. Electrode bursts are shown in blue and network bursts are outlined in pink. **H.** Northern blot for three mature miRNAs from miR-379/410 in adult mouse cortex and liver, *in vitro* ESCs, and iNs (day 4 and 8) relative to 5.8S rRNA.

**Figure S2.**
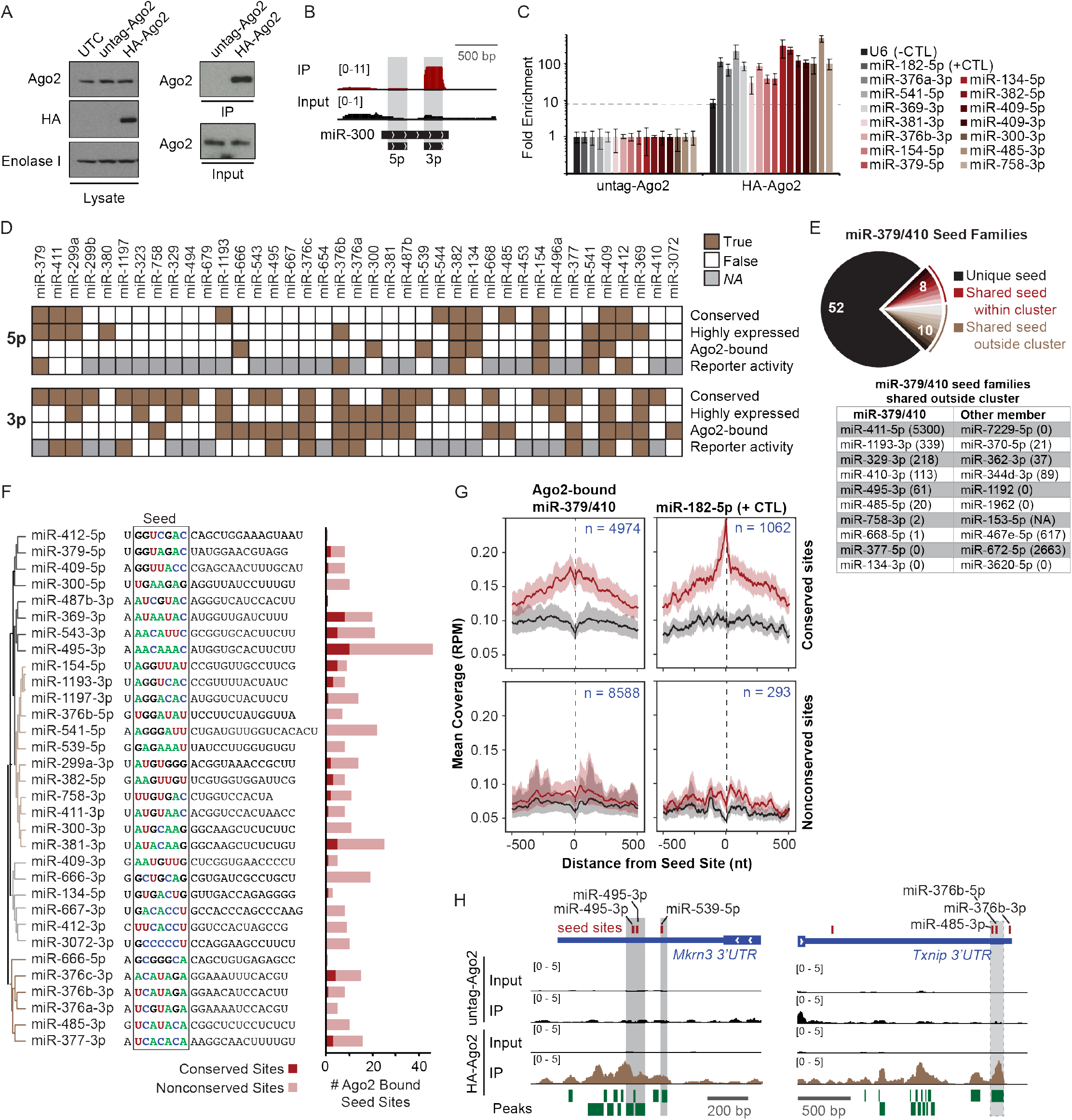
miR-379/410 targeting by Ago2 seCLIP in iNs. **A.** Western blot from cells with overexpression of untagged-Ago2 or HA-Ago2 from a stably integrated transgene. Left: whole cell lysates probed with antibodies for Ago2, HA, or Enolase I (loading control). UTC, untransfected control with endogenous Ago2 expression. Right: Confirmation of Ago2 immunoprecipitation in seCLIP-seq. Cross-linked cell lysates were immunoprecipitated with an HA antibody. The input and IP samples were then probed with Ago2 antibody. **B.** Genomic alignment of seCLIP-seq reads to miR-300. Scale is reads per million. **C.** RNA immunoprecipitation using an HA antibody for iNs expressing untagged-Ago2 or HA-Ago2 followed by miRNA qRT-PCR. Dotted line indicates enrichment threshold established by U6 snRNA negative control. miR-182-5p is a positive control from outside the miR-379/410 cluster. **D.** Summary of data for each miRNA from miR-379/410 organized according to their 5’-to-3’ position within the cluster. Conserved, true if miRNA is conserved among eutherian mammals (from TargetScanMouse, release 7.2); Highly expressed, from Figure 1B; Ago2-bound, from Figure 2A; Reporter activity, from Figure 1G. **E.** Top: Pie chart of seed families in miR-379/410 cluster. Unique seed families contain one miR-379/410 miRNA (n = 52, black), seed families with two miR-379/410 family members (n = 8, red), seed families with one miR-379/410 family member and one member outside the cluster (n = 10, gold). Bottom: Table of miR-379/410 seed families that share a seed sequence with a miRNA outside the cluster. Normalized expression counts in iNs for each miRNA are indicated in parenthesis. **F.** Active miR-379/410 miRNAs were clustered according to the similarity of their seed sequence. The mature miRNA sequence is shown and the seed sequence is highlighted. The bar graph indicates the number of significant Ago2 peaks that overlap a conserved or nonconserved seed match for each active miRNA. **G.** Metagene plot of Ago2 seCLIP-seq coverage (reads per million, RPM) at seed sites for the indicated miRNAs, including coverage 500 nt upstream and downstream of the seed site. The number of sites included in each plot is indicated. Top, conserved sites; Bottom, nonconserved sites with TargetScan score ≥ 90; Black, untagged-Ago2 IP; Red, HA-Ago2 IP. **H.** Genomic alignment of Ago2 seCLIP-seq reads to the 3’ UTR of *Mkrn3* and *Txnip*. Reads are scaled to reads per million (RPM). Predicted binding sites for active miR-379/410 miRNAs are indicated in red. Significant Ago2 peaks are shown as green boxes. Significant Ago2 peaks overlapping a seed site for miR-379/410 are highlighted in gray.

**Figure S3.**
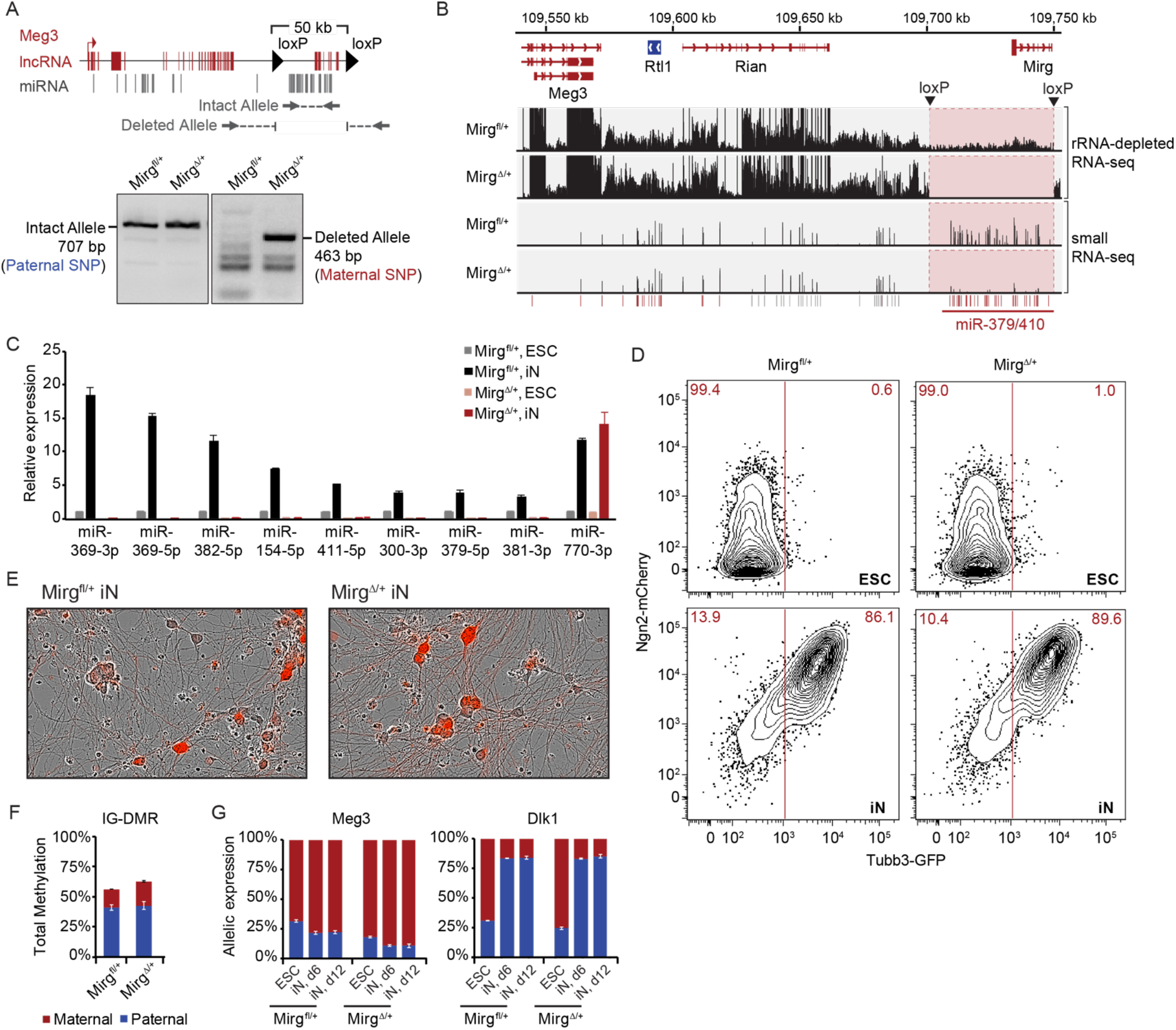
Characterization of maternal miR-379/410 deletion in ESCs. **A.** Top: Schematic of *Meg3* genomic locus showing location of maternal loxP insertions and primer locations to confirm presence of intact and deleted allele following Cre-mediated recombination. Bottom: Agarose gel of PCR products for intact and deleted allele in Mirg^fl/+^ and Mirg^Δ/+^ ESCs. Allelic identity was obtained by Sanger sequencing across allelic SNPs from gel isolated bands. **B.** Genomic alignment of rRNA-depleted RNA-seq and small RNA-seq reads to the Chr12 *Meg3* locus in Mirg^fl/+^ and Mirg^Δ/+^ iNs. **C.** qRT-PCR for mature miRNAs from miR-379/410 in Mirg^fl/+^ and Mirg^Δ/+^ ESCs and iNs. miR-770-3p is a miRNA transcribed outside the miR-379/410 cluster as an assay control. Mean ± SEM (n = 3). **D.** FACS of ESCs and iNs based on Ngn2-mCherry and endogenously tagged Tubb3-GFP. Percentage of Tubb3-GFP negative or positive cells is indicated. **E.** Overlay of brightfield and Ngn2-mCherry fluorescence images for Mirg^fl/+^ and Mirg^Δ/+^ iNs. **F.** Quantification of allele-specific DNA methylation at the intergenic differentially methylated region (IG-DMR) for the *Dlk1*/*Meg3* locus in Mirg^fl/+^ and Mirg^Δ/+^ ESCs using pyrosequencing. Mean ± SEM (n = 3). **G.** Allele-specific qRT-PCR for *Meg3* and *Dlk1* expression in Mirg^fl/+^ and Mirg^Δ/+^ ESCs and iNs. Mean ± SEM (n = 3).

**Figure S4.**
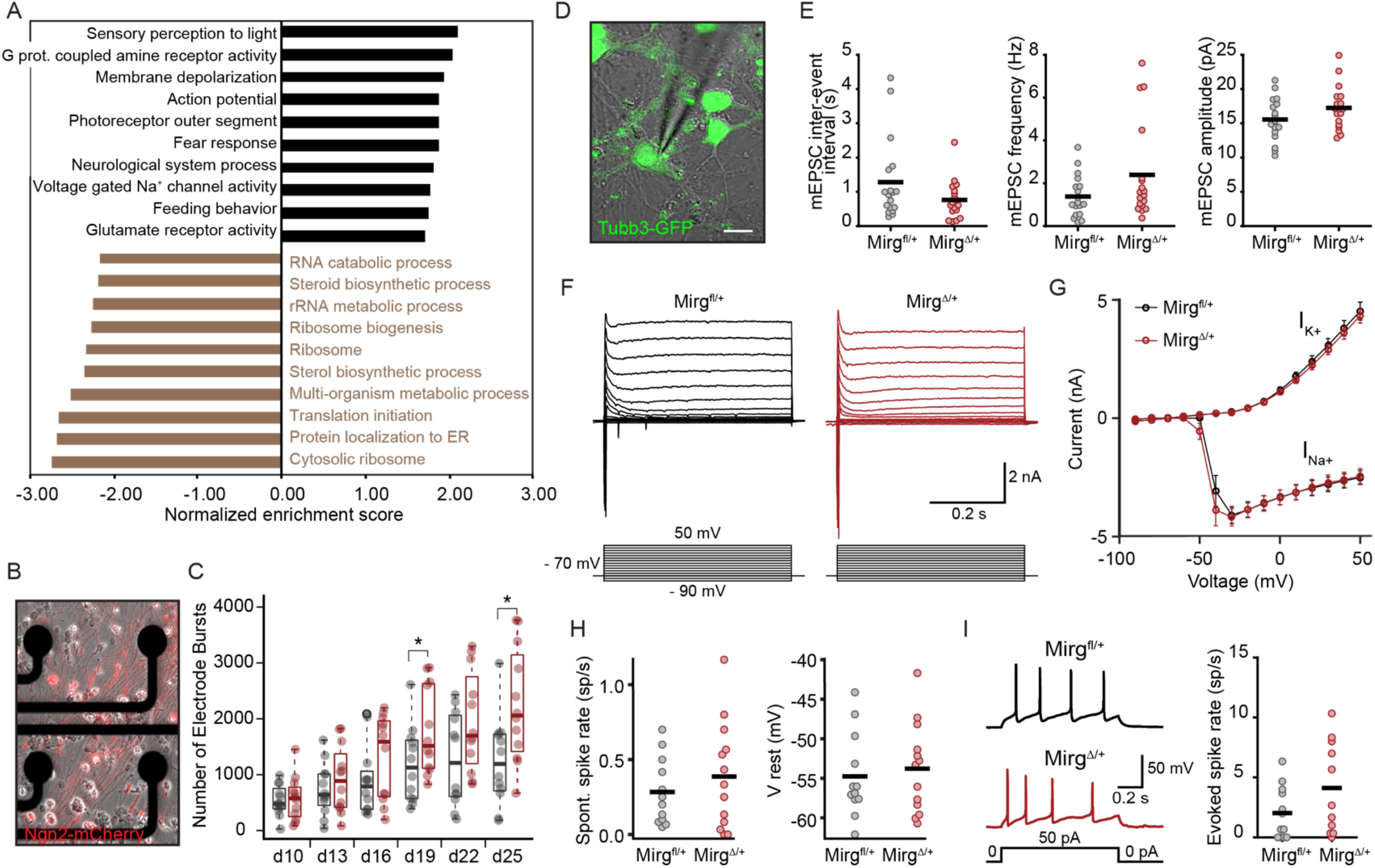
miR-379/410 deletion increases synaptic events but preserves spiking and membrane properties. **A.** Bar graph of normalized enrichment scores from gene set enrichment analysis for gene ontology terms associated with genes up-regulated (black) or down-regulated (gold) in Mirg^Δ/+^ iNs. **B.** Overlay of brightfield and Ngn2-mCherry fluorescence image of iNs growing on multielectrode array (MEA) plate. **C.** Boxplot of number of electrode bursts from MEA recordings at day 10 to 25 of differentiation. Individual data points are the mean of 16 electrodes in a single well. Black, Mirg^fl/+^ iNs (n = 12 wells); Red, Mirg^Δ/+^ iNs (n = 12 wells). **D**. Overlay of differential interference contrast (DIC) image and Tubb3-GFP fluorescence for one example recorded iN. Scale bar, 10 μm. **E**. Quantification of inter-event interval (p = 0.10, t_34_ = 1.6773), mEPSC frequency (p = 0.09, t_34_ = −1.7588), and amplitude (p = 0.11, t_34_ = 1.6237) from mEPSC voltage-clamp recordings. Individual data points are the mean values for each cell, n = 18 cells for each condition. Solid black bar represents the mean value for each condition. P-values were calculated using unpaired two-tailed Student’s t-test. **F**. Example voltage-clamp trace in response to voltage steps for Mirg^fl/+^ and Mirg^Δ/+^ iNs. **G**. Amplitude of sodium (I_Na+_) and potassium (I_K+_) currents at different voltage steps. Black, Mirg^fl/+^ iNs (n = 12); Red, Mirg^Δ/+^ iNs (n = 13); Mean ± SEM. **H**. Spontaneous spike rate (p = 0.40, t_23_ = −0.86) and resting membrane potential (p = 0.67, t_23_ = −0.44) measured during one-minute-long current-clamp recordings without current injection. Black, Mirg^fl/+^ iNs (n = 12); Red, Mirg^Δ/+^ iNs (n = 13). **I**. Left: Representative current-clamp traces of the spiking activity in response to +50 pA current steps for each condition. Right: Average evoked spike rate at 50 pA current injection (p = 0.12, t_21_ = −1.6335). Black, Mirg^fl/+^ iNs (n = 12); Red, Mirg^Δ/+^ iNs (n = 11). P-values in (**H**) and (**I**) were calculated using unpaired two-tailed Student’s t-test. Solid black bars in (**H**) and (**I**) represent the average value for each condition.

